# DrDimont: Explainable drug response prediction from differential analysis of multi-omics networks

**DOI:** 10.1101/2022.05.31.493964

**Authors:** Pauline Hiort, Julian Hugo, Justus Zeinert, Nataniel Müller, Spoorthi Kashyap, Jagath C. Rajapakse, Francisco Azuaje, Bernhard Y. Renard, Katharina Baum

**Affiliations:** Hasso Plattner Institute, Digital Engineering Faculty, University of Potsdam, Potsdam, 14482, Germany; School of Computer Science and Engineering, Nanyang Technological University, 639798, Singapore; Genomics England, London, EC1M 6BQ, United Kingdom

## Abstract

**Motivation:** While it has been well established that drugs affect and help patients differently, personalized drug response predictions remain challenging. Solutions based on single omics measurements have been proposed, and networks provide means to incorporate molecular interactions into reasoning. However, how to integrate the wealth of information contained in multiple omics layers still poses a complex problem.

**Results:** We present DrDimont, **D**rug **r**esponse prediction from **Di**fferential analysis of **m**ulti-**o**mics **n**e**t**works. It allows for comparative conclusions between two conditions and translates them into differential drug response predictions. DrDimont focuses on molecular interactions. It establishes condition-specific networks from correlation within an omics layer that are then reduced and combined into heterogeneous, multi-omics molecular networks. A novel semi-local, path-based integration step ensures integrative conclusions. Differential predictions are derived from comparing the condition-specific integrated networks. DrDimont’s predictions are explainable, i.e., molecular differences that are the source of high differential drug scores can be retrieved. We predict differential drug response in breast cancer using transcriptomics, proteomics, phosphosite, and metabolomics measurements and contrast estrogen receptor positive and receptor negative patients. DrDimont performs better than drug prediction based on differential protein expression or PageRank when evaluating it on ground truth data from cancer cell lines. We find proteomic and phosphosite layers to carry most information for distinguishing drug response.

**Availability:** DrDimont is available on CRAN: https://cran.r-project.org/package=DrDimont.

## 1 Introduction

Personalized prediction of suitable medication is still a key task for computational approaches in clinical research. Meta-studies have shown that many drugs only work effectively in a fraction of patients (Leucht *et al*., 2015), and consequences of failing treatment and adverse drug events can be severe. In recent years, more and more multi-omics profiles have become available that characterize disease phenotypes on a molecular level, especially in cancer, for example via the TCGA consortium (Chang *et al*., 2013). Multiple layers of molecular data provide different perspectives and a higher resolution. Thus, a more fine-grained distinction between subgroups of patients is possible. At the same time, these data present the challenge of how to integrate them in order to derive meaningful predictions.

Different methods of omics integration have been proposed, distinguished frequently by when the omics layers are combined as well as by the goal of the analysis – molecular mechanism, patient clustering, or other predictions such as drug response (Bersanelli *et al*., 2016; Huang *et al*., 2017; Picard *et al*., 2021). Thereby, a genuine joint integration of the different layers has been considered advantageous (Picard *et al*., 2021; Cantini *et al*., 2021). Joint dimensionality reduction, e.g., via ICA or MOFA (Sompairac *et al*., 2019; Argelaguet *et al*., 2018), has been proposed to find relevant features from combined multi-omics data. They have been benchmarked for use with cancer data (Cantini *et al*., 2021). However, the reduced meta-genes are difficult to interpret and hinder direct conclusions on drug action.

The fact that molecules do not act separately, but in their network context led to an alternative joint integration strategy: Network-based approaches enable consideration of interactions between entities and have been specifically applied to multi-omics data (Recanatini and Cabrelle, 2020; Yugi *et al*., 2016; Lee *et al*., 2019; Demirel *et al*., 2021). Networks other than purely molecular heterogeneous networks have been proposed, containing, e.g., diseases, drugs, or cell lines as nodes (Zhang *et al*., 2018a; Stanfield *et al*., 2017), and patient similarity networks derived from molecular data (Wang *et al*., 2014). A plethora of methods have been suggested to establish and use molecular networks to find relevant disease genes (Ogris *et al*., 2021; Dimitrakopoulos *et al*., 2018; Schulte-Sasse *et al*., 2021; Peng *et al*., 2017). Moreover, interactions between molecules are one of the key readouts of drug action: drugs interfere most frequently with the function of the targets they bind to, hampering their ability to interact with other molecular players instead of affecting their overall abundance (Pinto *et al*., 2014). Therefore, considering interactions in molecular networks (Bartel *et al*., 2015; Koh *et al*., 2019; Sambaturu *et al*., 2020) is highly promising for drug response prediction.

Systematic drug response measurements are available, especially for cancer cell lines (Rees *et al*., 2015; Yang *et al*., 2013; Barretina *et al*., 2012), and therefore, drug response prediction has been performed for this *in vitro* setting (Ding *et al*., 2018; Zhang *et al*., 2018a; Park *et al*., 2022). The question remains how these results can be transferred to clinically relevant predictions for patients. Some transfer learning approaches have been proposed (Geeleher *et al*., 2017; Webber *et al*., 2018), but these still rely on prior systematic measurements of drug response in conditions comparable to the condition of interest. Methods of artificial intelligence are being advanced (Azuaje, 2019), but the explanation of their results is frequently challenging. A viable strategy is to assess differential response to a baseline patient group phenotype. This technique has been proposed to query differential co-expression (Matsui *et al*., 2021; Bhuva *et al*., 2019), gene ranking (Richard *et al*., 2020), or networks or paths (Ideker and Krogan, 2012; Sambaturu *et al*., 2020), but not differential drug response yet.

Here, we present our new approach for **D**rug **r**esponse prediction from **Di**fferential analysis of **m**ulti-**o**mics **n**e**t**works, DrDimont, that unites the following key points: (i) multi-omics data is jointly integrated, including data such as metabolomics, (ii) condition-specific molecular networks are built, (iii) prior and domain-specific knowledge on molecular interactions can be leveraged, (iv) the focus is on interactions between molecules as the most common mode of drug action, (v) differential analysis between conditions enables unsupervised predictions in clinically relevant settings, and (vi) the predictions are explainable as their underlying molecular characteristics can be retrieved.

We describe DrDimont and apply it to a breast cancer dataset combining transcriptomics, proteomics, phosphosite, and metabolomics measurements. We compare DrDimont’s differential drug response predictions to ground truth from cell line measurements, and to alternative approaches. We investigate the impact of different measurement layers on the prediction quality. Finally, we showcase an example of how DrDimont explains results down to the molecular level.

## 2 Methods

### 2.1 Differential predictions with DrDimont

DrDimont provides a framework to leverage condition-specific, weighted heterogeneous networks for differential analysis between two conditions. It builds purely molecular networks with nodes encoding entities within a cell, such as proteins, mRNAs, metabolites, and their interactions from both multi-omics data and prior information on interactions from databases.

An overview of the pipeline provided in the DrDimont framework is shown in Figure 1. DrDimont requires quantified molecular data such as RNAseq or protein data of several samples as input. For differential analysis, data for two different groups of samples or patients (‘conditions’) are needed. Each molecular data input layer is transformed into condition-specific weighted, single-layer networks by correlation of the molecular entities. Then, based on a user-defined structural requirement (see *Single-layer network generation*), the networks are reduced keeping edges with high weights only. As shown for an example in Figure 1B, the single-layer networks are combined into multi-layer networks based on user-defined inter-layer connections. Thereby, prior knowledge on interactions from databases can be incorporated. The two condition-specific multi-layer networks are further integrated by computing integrated interaction scores, as shown in Figure 1C. These propagate local neighborhood information to the edge weights and thus avoid too strong of an impact of single edges. From the two integrated networks a differential network is computed by contrasting the edge weights of the condition-specific networks. The differential network is employed to calculate differential drug response scores based on the differential edges in vicinity to a drug’s targets. We will describe details for each step in the following.

**Fig. 1.**
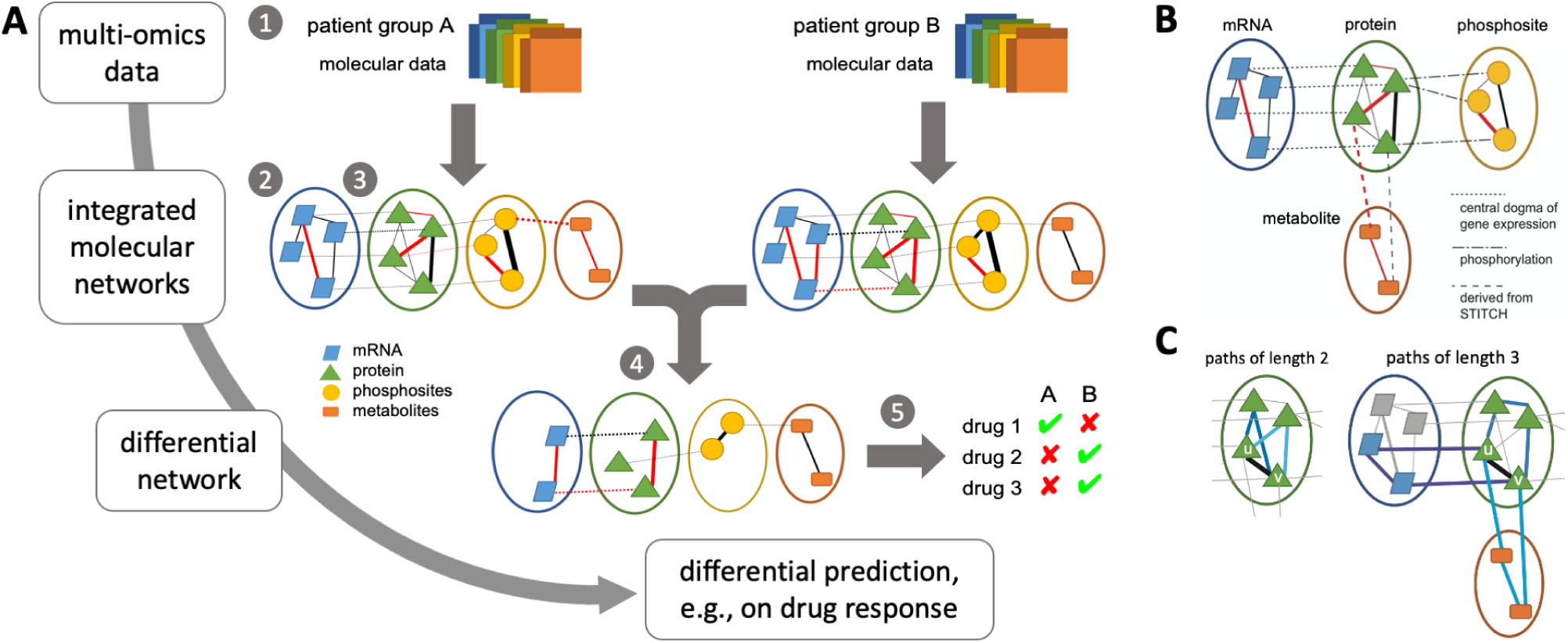
DrDimont’s pipeline for integrated, network-based analysis. (A) Multiple molecular data layers for two conditions are compared, e.g., cancer data (1). They are used to derive condition-specific correlation-based single-layer networks (2) that are combined into integrated molecular networks using prior information (3). The differential network is derived from the condition-specific networks (4) and captures altered interaction strengths. Differences, e.g., in drug response, are predicted from the differential network (5). (B) Example for the generation of an integrated, multi-layer weighted network with the protein network as central layer. Within-layer interactions are correlation-based from measurements. Different layers are connected by prior information, its type is encoded by the dashed lines. Red denotes negative edge weights, black positive edge weights, thicker lines indicate larger edge weights. (C) Integrated interaction scores. Edge weights are replaced by the average over the strengths of alternative paths, thus generating a local enhancement exploiting network structure (see Equation 1). For example, for the edge connecting nodes u and v, a selection of alternative paths is marked by blue edges: paths of length two (left), or paths of length three (right).

#### 2.1.1 Single-layer network generation

For each data layer and each group, complete weighted networks are generated where each node represents one type of molecules, e.g., one specific mRNA or protein. The weight of an edge between any two nodes of one layer is derived from the correlation between the abundance measurements for the nodes over all samples in the group, e.g., using Spearman’s or Pearson’s correlation. If not stated otherwise, we employed Spearman’s correlation in order to avoid strong impact of outliers and to account for non-linearity in abundance relationships. In case of missing values, pairwise complete observations were used for correlation.

The correlation-based networks are reduced and only the edges with the largest absolute weights are kept. Reduction thresholds are determined (i) by a desired average degree of the network nodes, (ii) by a desired average network density, or (iii) by maximizing the scale-freeness of the network (WGCNA, pickHardthreshold function (Langfelder and Horvath, 2008)). If not stated otherwise, we used this latter topological criterion here and adapted the goodness of fit to a scale-free network, R^2^, to have similar-sized networks for the compared groups. See the Supplement for the impact of *Alternative reduction methods*.

#### 2.1.2 Heterogeneous multi-layer network construction

DrDimont connects single-layer networks for each group separately based on the node names (see Figure 1B). First, nodes from different layers with identical names can be connected with edges of the same user-defined weight (e.g., of value one). This allows exploiting the dogma of gene expression, i.e., connecting an mRNA to its corresponding protein, or a protein to its corresponding phosphosites. Representing further relationships is possible such as methylated promoter regions on the DNA to the corresponding target genes or mRNAs. Second, pairs of node names from two different layers can be entered by the user. These nodes will then be connected. The edge weights can be again fixed or derived from prior information, for example, to connect the protein and the metabolite layer using data from a database such as STITCH (Szklarczyk *et al*., 2016) (see Supplement, *Metabolite-protein interactions from STITCH* for details).

#### 2.1.3 Integrated interaction scores

DrDimont uses a novel, semi-local integration scheme to reduce the impact of single edge weights in the condition-specific networks. Thereby, the weights of alternative paths between nodes are taken into account (see Figure 1C). Edge weights in the heterogeneous multi-layer network can be replaced by their integrated interaction scores, that is, the average strength of these alternative paths. For an edge connecting nodes *u, v*, we define this score, *s*_*u,v*_, as the sum of average strengths of alternative paths connecting *u* and *v* over the considered path lengths, i.e.,

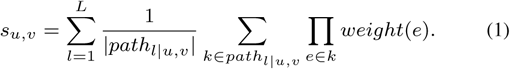

Thereby, *path*_*l*|*u,v*_ is the set of paths of length *l* between nodes *u* and *v, L* is the maximal length of considered paths, and *weight*(*e*) is the edge weight of an edge *e*. For a path *k* of length *l* that is connecting nodes *x*_0_, …, *x*_*l*_ in this order, *k* is determined by its set of contributing edges *k* = {(*x*_0_, *x*_1_), (*x*_1_, *x*_2_), …(*x*_*l*−1_, *x*_*l*_)}, and all their edge weights are multiplied to determine the path strength of *k*. For edge weights ranging from −1 to 1 as in usual correlation measures, integrated interaction scores can range from −*L* to *L*. If not stated otherwise, we used *L* = 3. In order to reduce DrDimont’s run time for large networks, the integrated interaction scores are only computed for edges incident to user-provided drug targets by default (see also *Differential drug response score*).

#### 2.1.4 Differential network

DrDimont generates the differential network by computing the difference between integrated interaction scores of all edges of the group-specific heterogeneous multi-layer networks. Edges that only appear in one of the group’s networks are considered to have a weight of zero in the other networks and will be part of the differential network. Nodes that are part of any of the two networks are thus included in the differential network.

#### 2.1.5 Differential drug response score

The differential network is used for DrDimont’s differential drug response prediction. To derive a prediction for a drug, drug targets have to be known. The drug targets are identified within the differential network. The absolute value of the mean (default, or the median) of the weights of all edges incident to the drug targets is used as a drug’s differential drug response score. The differential drug response score is the main output of DrDimont and provides a prioritization (ranking) of the drugs. If not stated otherwise, we used proteins as drug targets. However, in principle, nodes from any input molecular layer can be defined as drug targets.

DrDimont’s implementation details and its settings for heterogeneous network construction are provided in the Supplement.

### 2.2 Molecular breast cancer dataset preparation

We used a breast cancer dataset from patient tumors with mRNA (measured via RNAseq) from TCGA, proteomics and phosphosites (measured via mass spectrometry) from CPTAC (Mertins *et al*., 2016). We combined these data with metabolite data from two other studies (Budczies *et al*., 2013; Terunuma *et al*., 2014). The mRNA data and clinical annotations were downloaded from TCGA via RTCGA (Kosinski and Biecek, 2021). We obtained the estrogen receptor status, negative (ER-) or positive (ER+), for each sample from the clinical annotations (for sample counts, see Supplement, Table S1).

We disregarded mRNAs with more than 90% of zero measurements over the samples within a condition. Proteins and phosphosites with more than 20% of missing values over the samples of a condition were removed. There were no missing values for the metabolite data as imputation had been done by the respective authors prior to publication of the data. If not stated otherwise, the genetic features (mRNA, protein, phosphosites) were reduced to a subset of 5579 known cancer-related genes (Repana *et al*., 2019) and drug targets from DrugBank (Wishart *et al*., 2017).

### 2.3 Drug targets, drug response ground truth, performance

We retrieved data for 40 breast cancer cell lines (26 ER-, 14 ER+) from the Cancer Therapeutics Response Portal (CTRP) (Rees *et al*., 2015), in particular drug sensitivity, compound data and drug target information for 481 drugs. We used estrogen receptor status annotation from the DepMap portal (DepMap, Broad, 2021). We employed data for a drug if it was measured at least three times for each condition. We determined the differential drug response between ER+ and ER-for each drug by Mann-Whitney-U tests comparing sensitivity in ER+ cell lines vs. sensitivity in ER-cell lines obtaining ground truth for 477 drugs with the p-value as ranking (see Supplement, Figure S1, for the effect size instead of p-value). For performance assessment, we report Spearman’s correlation between predicted and ground truth drug ranking, and the p-value of the correlation. Only drugs for that the analysis in question delivered a prediction were used; their numbers are indicated accordingly. In particular, drugs lacking known drug targets were disregarded.

Receiver operating characteristic (ROC) curves were generated by comparing prediction-derived to ground-truth derived binary drug classifications (see Supplement for details). We indicate which fixed ground-truth threshold was employed. We report the area under the ROC curve (AUC). For partial AUC (pAUC), we compute the AUC for false positive rates between 0 and 0.1. High pAUC values signify enriched true predictions among the top ranked drugs.

Data preparation, visualization and result analysis was performed using R, version 4.0.1 (R Core Team, 2021).

### 2.4 Alternative prediction methods

To assess DrDimont’s performance, we implemented two alternative differential drug response prediction approaches.

#### 2.4.1 PageRank of drug targets in the differential network

We used igraph (Csardi and Nepusz, 2006) to compute the weighted PageRank (Brin and Page, 1998) of all potential drug targets in the undirected differential network from DrDimont. Therein, we employed the absolute differential edge weights (and not differential integrated interactions scores). The mean weighted PageRank over all drug targets generated the differential drug response ranking.

#### 2.4.2 Differential protein expression

We employed the Differential Enrichment analysis of Proteomics data (DEP) pipeline (Zhang *et al*., 2018b) to assess differential protein expression between ER+ and ER-conditions. Therein, after a variance-stabilizing transformation, *limma* is applied. This single-layer approach delivered a ranking of the drugs with respect to differential drug prediction from differential protein expression of all drug targets of a drug using the minimal multiple-testing adjusted p-value.

## 3 Results

We will now showcase results of DrDimont for a multi-omics dataset and assess its performance with measured ground truth. Then, we will compare it to two alternative differential drug response prediction approaches. Furthermore, we will describe the impact of including different data layers in DrDimont’s analysis, and end with an illustration of the level of explainability that DrDimont provides.

### 3.1 Evaluation of DrDimont on breast cancer stratified by estrogen receptor status

Breast cancer is one of the most common cancers. We investigated molecular data of breast cancer patients stratified by estrogen receptor status. The ER status is highly prognostic, with ER+ patients having a better prognosis than ER-patients.

We first considered a multi-omics dataset that provides ER-stratified patient data containing transcriptomics measurements via RNAseq (from TCGA), and proteomics and phosphosite data from mass spectrometry-based measurements (from CPTAC, (Mertins *et al*., 2016)). We also included metabolomics data from other studies later (Budczies *et al*., 2013; Terunuma *et al*., 2014). We show the properties of the differential integrated network generated with DrDimont for this dataset (without metabolomics) in Figure 2A. The differential integrated interaction scores are correlated with the differential edge weights, but the former allow for broader distributions. They are taking alternative paths into account and thereby propagate the information from the local neighborhood to the respective edges. In particular, the edges derived from prior information (mRNA-protein, protein-phosphosite) benefit from this procedure: Their edge weights are not condition-specific and therefore, their differential edge weights cluster around minus one, zero, and one. In contrast, their differential integrated interaction scores are spread out and differences between conditions are resolved in greater detail.

**Fig. 2.**
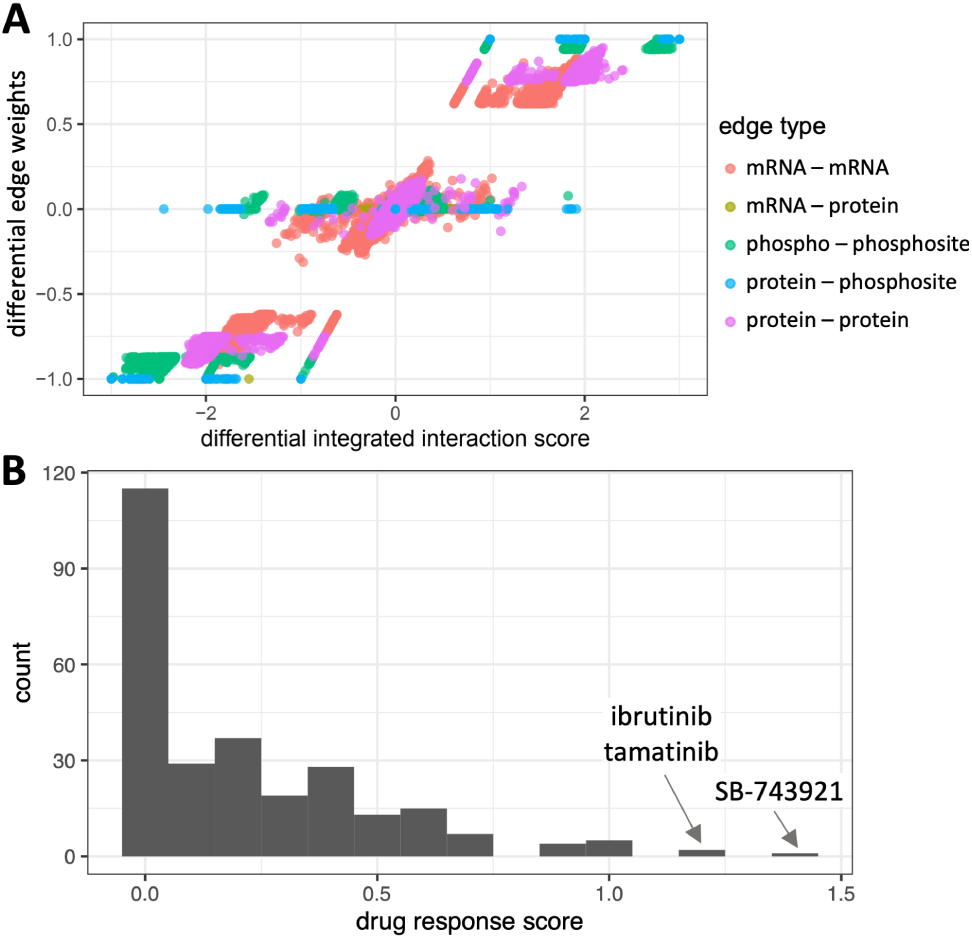
DrDimont’s integrated interaction scores and differential drug response predictions for the breast cancer dataset. (A) Differential edge weights are compared to the differential integrated interaction scores of DrDimont for each of the five edge types (colored). We contrasted ER-vs. ER+, i.e., negative scores correspond to stronger inhibitory interactions or less strong positive interactions in ER-compared to ER+. Integrated interaction scores enable distinguishing edges that have a zero differential edge weight, but overall, they show a high correlation (Spearman’s *ρ* 0.967). (B) Histogram of DrDimont’s differential drug response scores of 275 drugs for which drug targets occur in our multi-layer networks. Half of the drugs are predicted with a differential response of varying value.

For our dataset, DrDimont provides drug response scores for 275 drugs with drug targets from CTRP (Rees *et al*., 2015) (see Figure 2B), the majority were differentially predicted to some degree. Only seven drugs have a drug response score of one or higher. Top differentially predicted drugs were SB-743921, that is known to have a stronger effect in ER-cell lines (Zhu *et al*., 2016), ibrutinib, and tamatinib.

We compared the results of DrDimont for our dataset to the ground truth of the drugs from CTRP in a receiver operating characteristic (ROC) performance analysis (see Figure 3A). DrDimont’s drug response scores were used as drug ranking for computation of true and false positive rates of the prediction. The area under the ROC curve (AUC) for a ground truth threshold of 0.01 was 0.67 (see legend of Figure 3A) which is considerably higher than for random prediction (AUC 0.5). DrDimont’s multi-omics data-based drug response scores showed a significant correlation to the CTRP-based ground truth (Spearman’s *ρ* -0.19, p-value 0.001). The correlation is negative since the ground truth is based on p-values, i.e., lower values correspond to a likelier differential response. What is more, the partial AUC (pAUC) that takes only highly ranked drugs into account gives values decisively higher than expected from a random prediction (up to 0.014 compared to 0.005). Thus, we find DrDimont to be predictive of differential drug response, and it especially enriches positive hits among the top ranked results.

**Fig. 3.**
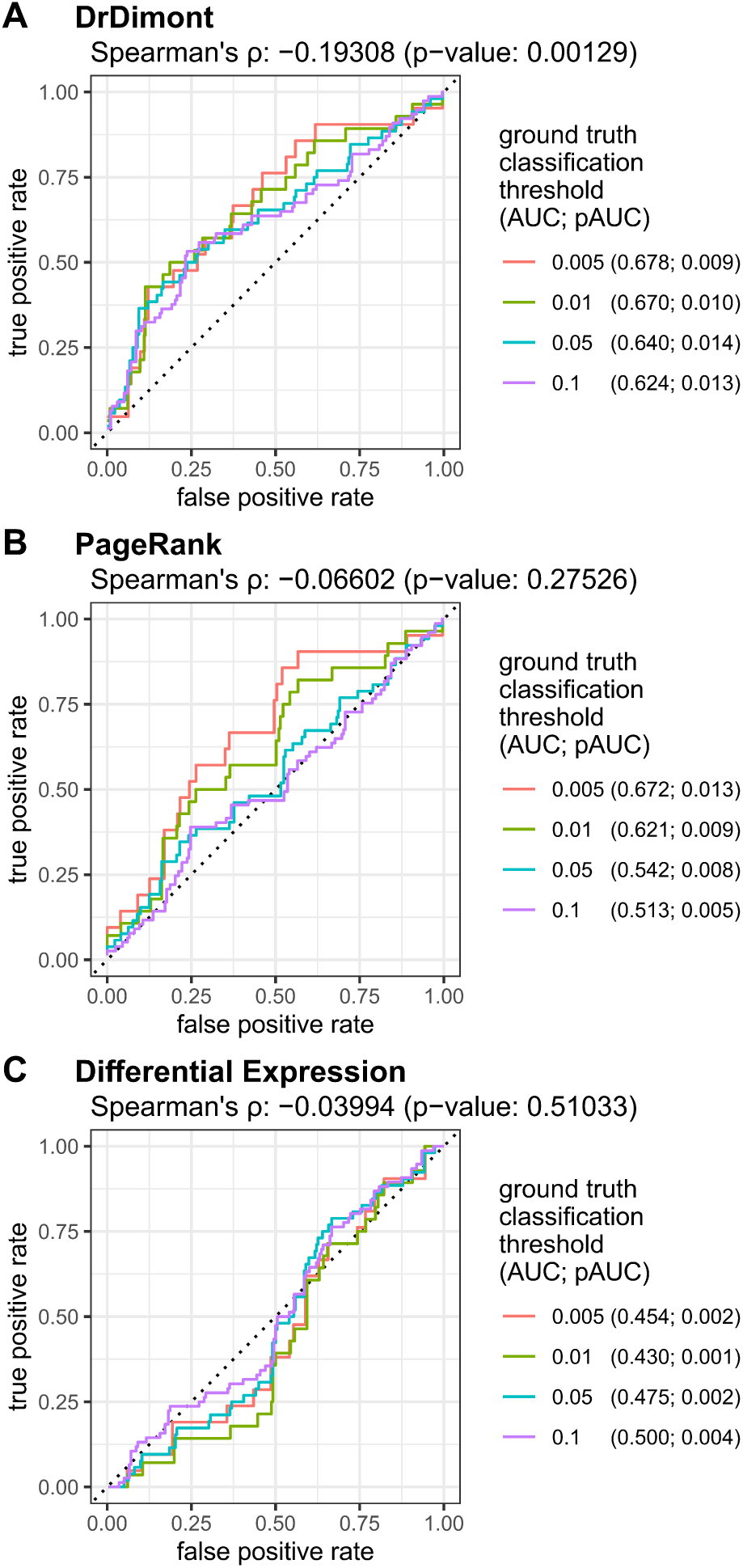
ROC curves for DrDimont’s and alternative methods’ prediction performance on the breast cancer dataset. (A) DrDimont’s differential drug response. Different ground truth thresholds were used where drugs below the threshold were considered as true positives and the others as true negatives. For 275 drugs, DrDimont provided differential drug response scores that were used for drug ranking and comparison to ground truth classification. (B) Weighted PageRank of drug targets for differential drug response prediction, for 275 drugs, computed on DrDimont’s differential network. (C) Differential expression of drug target proteins for differential drug response prediction. The differential expression yielded predictions for 274 drugs. In the legends, values in brackets denote AUCs, pAUCs at given ground truth thresholds. The respective Spearman’s *ρ* and the p-value are given at the top of each figure. Black dotted lines indicate theoretical ROC curves for random predictions.

### 3.2 Comparison to alternative differential drug prediction approaches

DrDimont’s approach focuses on differential interactions of drug targets in the molecular network for differential drug response predictions. However, frequently, rather the node properties are considered for predictions. Therefore, we compare DrDimont’s performance to two alternative approaches for deriving differential drug response: (i) using the weighted PageRank algorithm in DrDimont’s differential network to score drug targets, and (ii) differential expression of drug targets. PageRank detects nodes with high importance in a network (hubs) and could thus identify drug targets that are particularly altered between conditions from the differential network. Differential expression has been considered relevant for drug action and has been used especially for predictions of drug responsiveness of different tissues. We based our estimations on the differential protein expression of a drug’s targets because most frequently these are the drug’s binding partners rather than, e.g., mRNA molecules.

For our breast cancer dataset, the weighted PageRank for drug targets yielded predictions better than random, especially for stringent ground-truth thresholds (AUCs > 0.5, Figure 3B). Spearman’s correlation with the ground-truth ranking was less pronounced than for predictions with DrDimont and not significant (-0.06, p-value 0.27), and pAUCs were below random (pAUC < 0.005). The predictions based on differential protein expression were close to random classification (AUC ≤ 0.5). Also the correlation between prediction and ground truth ranking was small and insignificant (Spearman’s *ρ* -0.03, p-value 0.5, see Figure 3C). Thus, both the PageRank-based and the differential expression-based drug response predictions performed worse than DrDimont on the breast cancer dataset.

### 3.3 Influence of different molecular layers

We analyzed which molecular layers are most relevant for DrDimont’s drug response prediction performance. Therefore, we considered mRNA, protein, phosphosite data as before, as well as the two metabolomics datasets (see Table 1, and Supplement, Figure S4, for the ROC curves). DrDimont performed best according to AUC and Spearman’s correlation when employing only the proteomic layer in the analysis. However, DrDimont could only provide predictions for 116 drugs in this setting; the remaining drugs lacked edges for all of their drug targets in the network thereby resulting in no drug response score (see Supplement, Figure S5, for a further characterization). Compared to the default setting with all three data layers (no metabolomics), using protein and phosphosite data together resulted in only slight changes in AUC and correlation, but more than doubled the pAUC from 0.01 to 0.024 (best value). DrDimont performed worst in terms of AUC and correlation when using only the mRNA and the protein layers together. Applying DrDimont using only the phosphosite layer resulted in a relatively high pAUC of 0.017. Surprisingly, adding the two different metabolite datasets to the analysis with DrDimont showed consistently worse results than without metabolites, but affected the results for the setting containing all other three data layers least. We conclude that the proteomic and phosphosite data layers contribute most to DrDimont’s differential drug response prediction for this dataset.

**Table 1.** Influence of different molecular data layers on DrDimont’s drug response prediction performance. T: Terunuma metabolomics, B: Budczies metabolomics. The respective number of drugs with differential drug response predictions can change between approaches because the established molecular networks differ and drugs without any edges incident to their drug targets do not receive a differential drug response score. We indicate the AUC and pAUC for a ground truth threshold of 0.01, and Spearman’s correlation (corr.) and its p-value (p).

| included layers | #drugs | AUC | pAUC | corr. | p |
| --- | --- | --- | --- | --- | --- |
| mRNA, proteins, phosphosites | 275 | 0.670 | 0.010 | -0.193 | **0.001** |
| + metabolites B | 275 | 0.634 | 0.009 | -0.179 | 0.003 |
| + metabolites T | 275 | 0.634 | 0.007 | -0.172 | 0.004 |
| mRNA, proteins | 275 | 0.552 | 0.008 | -0.058 | 0.338 |
| mRNA | 216 | 0.574 | 0.001 | -0.114 | 0.093 |
| proteins, phosphosites | 190 | 0.668 | **0.024** | -0.188 | 0.009 |
| + metabolites B | 213 | 0.626 | 0.008 | -0.156 | 0.023 |
| + metabolites T | 194 | 0.643 | 0.007 | -0.145 | 0.034 |
| proteins | 116 | **0.677** | 0.021 | **-0.253** | 0.006 |
| + metabolites B | 178 | 0.479 | 0.008 | -0.035 | 0.639 |
| + metabolites T | 136 | 0.509 | 0.008 | -0.058 | 0.439 |
| phosphosites | 115 | 0.585 | 0.017 | -0.179 | 0.055 |

### 3.4 Explainable results

An asset of DrDimont is that predictions can be directly associated to molecular differences between subgroups. Given a specific drug response score for a certain drug, it can be traced back which drug target is especially different between the compared conditions, as well as which connections of the drug targets are the cause of the observed differences. We show this in an example for the drug dinaciclib, see Figure 4. Dinaciclib’s four reported drug targets CDK1, CDK2, CDK5, CDK9 and their incident edges can be identified both in the differential network as well as in the network for each condition. We find that CDK2 (and CDK5, not shown) has stronger interactions with other proteins in ER-than in ER+, whereas the interactions between proteins and their phosphosites are equally strong or stronger for CDK1 and CDK2 in the ER+ group. CDK9 shows no interaction differences between conditions. Specific differently interacting proteins and phosphosites can be also retrieved. This allows a deeper investigation by domain experts.

**Fig. 4.**
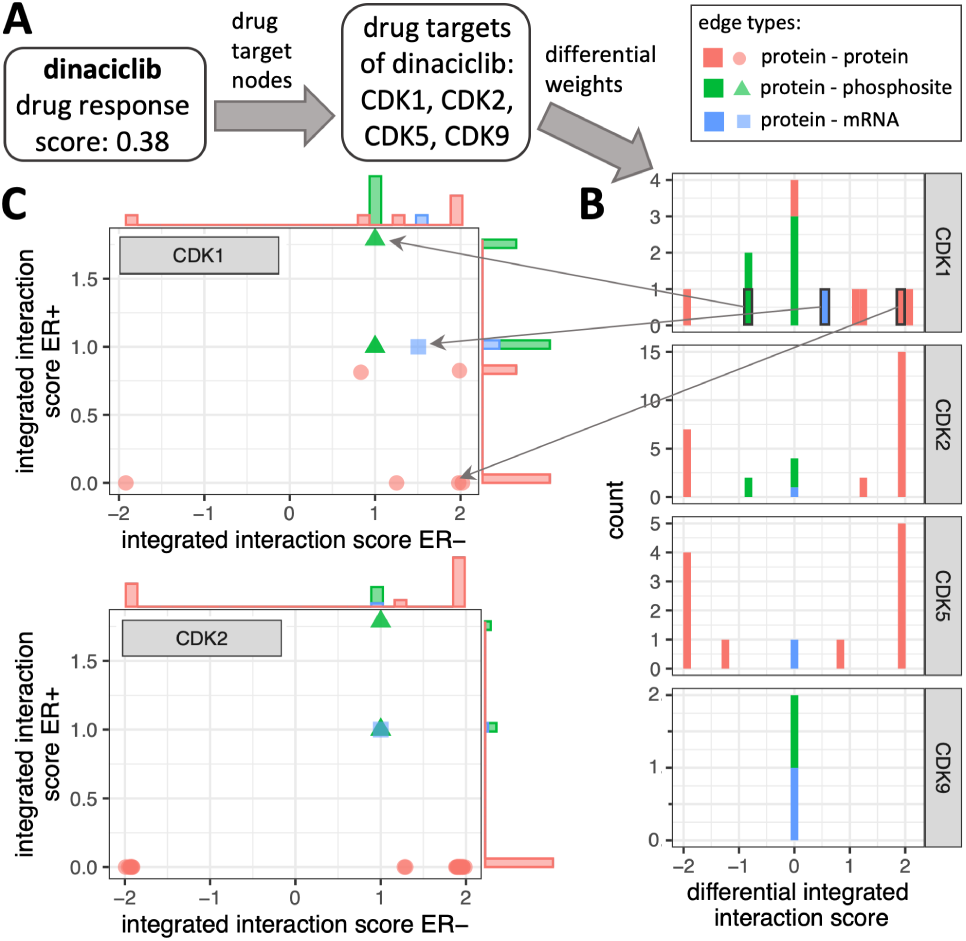
DrDimont delivers explainable predictions. Given a drug (example: dinaciclib), the differential drug response score derived from DrDimont can be traced back to the input layers. (A) Dinaciclib’s drug targets (CDK1, CDK2, CDK5, CDK9) are identified in the differential network. (B) The distribution of differential integrated interaction scores of the drug targets’ incident edges can be retrieved (stacked histogram). Many edges that are differential occur for CDK2 and CDK5. (C) The differential integrated interaction score of an edge can be related to the interaction strength in each condition (here: ER+ vs. ER-, for CDK1 and CDK2). Bars to the sides resolve edge counts. Boxes and arrows mark respective values in (B) and (C) for three selected edges incident to CDK1. Different edge types are marked by colors.

## 4 Discussion

We introduce DrDimont as a flexible framework for network-based differential analysis and drug response prediction. It builds condition-specific molecular networks from multi-or single-omics, and no matched samples are required. In addition, DrDimont outperforms differential expression-based and PageRank-based approaches for drug response prediction. It provides an explainable framework to trace contributions of single molecular alterations.

We find the protein and phosphosite layers to be most informative for drug prediction, especially for identifying top ranked drugs. Biologically, this seems reasonable because drugs mainly act on proteins where they interfere with their cellular functions (Pinto *et al*., 2014). Further, these functions are frequently modulated by post-translational modifications such as phosphorylations (Dittmar *et al*., 2019). Despite insights on the relevance of metabolomics in combination with other omics data for disease and in particular for cancer (Ortmayr *et al*., 2019; Pinu *et al*., 2019), we do not find evidence supporting that in our analyses. A possible reason could be that while all other omics data are from the same study and experimental conditions, the metabolomics measurements stem from different studies possibly adding extra noise to the integrated results. In addition, reliable measurement of metabolite abundances is more difficult in a patient treatment setting because metabolites are degraded extremely rapidly also within extracted tumor biopsies. Besides improvements in data quality and matched samples, further improvement may be achieved by reducing the number of the other layers’ features (nodes) for better balance of their sizes. Also alternative connection approaches might be a viable option to explore this further within the flexible framework of DrDimont. DrDimont’s combined molecular networks can be retrieved and employed for the user’s own analysis approaches, for example using network embedding (Pio-Lopez *et al*., 2021), exploration by random walks (Valdeolivas *et al*., 2019), or diffusion-based methods (Di Nanni *et al*., 2020). A strength of DrDimont is that its networks can also be tracked to provide molecular explanations for the predictions and thereby enable targeted biological follow-ups for in-depth investigation. In addition, DrDimont allows the inclusion of molecules with unknown function and low abundances. This makes the approach particularly interesting also for less well characterized organisms such as fungi.

Applying DrDimont to compare more refined subgroups would be interesting, for example, resolving effects of other hormone receptors in breast cancer such as progesterone or HER2. Other groups to compare could be pre-and post-chemotherapeutic patients (Park *et al*., 2020), or applying general sample classification before analysis with DrDimont.

A limitation of our network-based drug response prediction is that it relies on the quality of known drug targets. Some drugs are less well characterized than others. Taking this uncertainty into account and increasing the resolution on the mode of action of drugs for the input data (Parvizi *et al*., 2020) could be opportunities for improving DrDimont’s prediction results. Moreover, real ground truth for our case is difficult to obtain: applying different drugs to patients and monitoring their outcome cannot be performed, of course, due to ethical reasons and standard of care. Here, we use cell line measurements as a surrogate that differ in quality, taking trade-offs into account. Overall, differential drug response prediction is a difficult problem that also results in contradicting ground truth measurements (see Supplement, Figure S6), and thus, comparably low AUCs are not surprising. The partial AUC that measures enrichment of correct predictions in top-differential drugs achieves high values. Other directions such as relying on extended patient-derived xenograft-based drug response studies and analyses (Gao *et al*., 2015) could be taken.

DrDimont is a flexible tool for subgroup-specific and comparative predictions with an explainable framework, and we envision that it contributes with its proof-of-principle to improving the clinical decision process in the future.

## Acknowledgements

We thank Tim Garrels for assistance with implementation. The results shown here are in part based upon data generated by the TCGA Research Network: https://www.cancer.gov/tcga. Data used in this publication were generated by the Clinical Proteomic Tumor Analysis Consortium (NCI/NIH).

## Supplementary Information

### 1 Details on methods and settings

For general data analysis and visualization, we used R, version 4.1.0 (R Core Team, 2021). In addition, we employed the R packages ggplot2 (Wickham, 2016), ggextra (Attali and Baker, 2022), dplyr (Wickham *et al*., 2021), tidyr (Wickham and Girlich, 2022), and Hmisc (Harrell Jr, 2021).

#### 1.1 DrDimont’s implementation details

DrDimont is implemented as an R package and employs python (Python Core Team, 2021). The package was tested with R version 4.1.3 and python 3.9.6. R packages employed by DrDimont include igraph (Csardi and Nepusz, 2006), WGCNA (Langfelder and Horvath, 2008), dplyr (Wickham *et al*., 2021), tibble (Müller and Wickham, 2021), tidyr (Wickham and Girlich, 2022), stringr (Wickham, 2019), Rfast (Papadakis *et al*., 2022), readr (Wickham *et al*., 2022), magrittr (Bache and Wickham, 2022), rlang (Henry and Wickham, 2022), rmarkdown (Allaire *et al*., 2022), knitr (Xie, 2022), utils, and stats. Python packages used are numpy (version 1.20.3) (Harris *et al*., 2020), tqdm (da Costa-Luis *et al*., 2021), python-igraph (version 0.9.9) (Csardi and Nepusz, 2006) and ray (version 1.6.0) (Moritz *et al*., 2018).

#### 1.2 DrDimont’s settings for heterogeneous network construction

For the analysis of our datasets, DrDimont combined the mRNA, protein and phosphosite layers based on the gene names using edge weights of 1, i.e., an mRNA was linked to its protein, and a protein was linked to its phosphosites. The metabolites were linked to the proteins employing STITCH (Szklarczyk *et al*., 2016) interactions with confidence scores as edge weights (see *Metabolite-protein interactions from STITCH*).

#### 1.3 Metabolite-protein interactions from STITCH

Since combining the metabolite and protein or other genetic layers is not as straightforward, we downloaded all chemical-protein interactions from STITCH (Szklarczyk *et al*., 2016) for *Homo sapiens*. We restricted them to high-confidence interactions with a STITCH combined score > 900 and the sum of STITCH sub-scores for database and experimental evidence > 800. Interaction weights between proteins and metabolites were set as the combined STITCH score divided by 1000, and interactions were assigned a negative weight if they were identified as inhibition. We identified an interaction as inhibition if high-confidence evidence (score > 800) was given in the STITCH table of mode of actions. If both activation and inhibition were indicated for an interaction, we used the mode with a higher confidence. For the proteins, we mapped the STITCH ensembl_peptide_id to gene symbols using biomaRt. We used the aliases file from STITCH for identifying metabolites in our datasets based on Pubchem IDs, ChEBI IDs, KEGG IDs and metabolite names (in that order). The thus identified stereo chemical IDs for the aliases were used to match the chemical IDs of the STITCH interactions to the metabolite names given in the case study datasets.

#### 1.4 Performance assessment via ROC, AUC, pAUC

For receiver operating characteristic (ROC) analysis, a threshold for the ground truth ranking was applied to define a binary classification of drugs that have a differential response (low p-value, true positives) or have no differential response (high p-value, true negatives) in the ER-vs. ER+ cell lines. For the ROC curves of a predicted drug ranking, thresholds were varied to obtain predicted binary drug classifications. These were compared with the ground truth binary classification to derive false positive and true positive rates. The area under the ROC curve (AUC) was computed by numerical integration. The partial AUC (pAUC) was computed as area under the ROC curve between false positive rates of 0 and 0.1. Thus, random predictions yield pAUCs of 0.005. High pAUC values signify enriched true predictions among the top ranked drugs.

#### 1.5 Sample counts, network sizes, reduction thresholds

**Table S1.** Number of samples, nodes, reduced edges and DrDimont settings for network reduction per dataset, group and molecular layer. The network reduction parameters used in DrDimont were applied to both groups that we compared respectively. ‘red. TCGA’ denotes the TCGA + CPTAC dataset with reduced genetic features (our default) and ‘TCGA’ the unreduced dataset. The other two datasets, Krug (Krug et al., 2020) and cell lines (Li et al., 2019; Nusinow et al., 2020; Iorio et al., 2016), are described further in section Performance for unreduced and other datasets. The datasets Budczies (Budczies et al., 2013) and Terunuma (Terunuma et al., 2014) are the metabolomics datasets.

|  | Group | red. TCGA | TCGA | Krug | Cell lines | Budczies | Terunuma |
| --- | --- | --- | --- | --- | --- | --- | --- |
| mRNA |  |  |  |  |  |  |  |
| samples | ER- | 237 | 237 | 34 | 38 | - | - |
| samples | ER+ | 806 | 806 | 78 | 20 | - | - |
| nodes | ER- | 4120 | 18321 | 21015 | 18272 | - | - |
| nodes | ER+ | 4120 | 18321 | 20747 | 17938 | - | - |
| reduced edges | ER- | 21630 | 262787 | 86283 | 61703 | - | - |
| reduced edges | ER+ | 25036 | 138134 | 30498 | 34941 | - | - |
| R<sup>2</sup> | both | 0.85 | 0.85 | 0.87 | 0.89 | - | - |
| WGCNA threshold range | both | 0.4-0.8 | 0.55-0.8 | 0.56-0.8 | 0.55-0.9 | - | - |
| protein |  |  |  |  |  |  |  |
| samples | ER- | 36 | 36 | 34 | 23 | - | - |
| samples | ER+ | 68 | 68 | 78 | 9 | - | - |
| nodes | ER- | 2917 | 8430 | 9018 | 7443 | - | - |
| nodes | ER+ | 2920 | 8459 | 9011 | 7157 | - | - |
| reduced edges | ER- | 17634 | 81777 | 40621 | 83959 | - | - |
| reduced edges | ER+ | 6073 | 151712 | 55639 | 74741 | - | - |
| R<sup>2</sup> | both | 0.84 | 0.84 | 0.86 | 0.85 | - | - |
| WGCNA threshold range | both | 0.7-0.8 | 0.7-0.85 | 0.65-0.8 | 0.7-0.95 | - | - |
| phosphosites |  |  |  |  |  |  |  |
| samples | ER- | 36 | 36 | 34 | - | - | - |
| samples | ER+ | 67 | 67 | 78 | - | - | - |
| nodes | ER- | 4942 | 12290 | 18222 | - | - | - |
| nodes | ER+ | 4942 | 12223 | 17885 | - | - | - |
| reduced edges | ER- | 15109 | 34377 | 57845 | - | - | - |
| reduced edges | ER+ | 80674 | 10960 | 23468 | - | - | - |
| R<sup>2</sup> | both | 0.95 | 0.94 | 0.97 | - | - | - |
| WGCNA threshold range | both | 0.7-0.95 | 0.85-0.95 | 0.7-0.9 | - | - | - |
| metabolites |  |  |  |  |  |  |  |
| samples | ER- | - | - | - | 30 | 41 | 34 |
| samples | ER+ | - | - | - | 18 | 143 | 33 |
| nodes | ER- | - | - | - | 225 | 162 | 342 |
| nodes | ER+ | - | - | - | 225 | 162 | 307 |
| reduced edges | ER- | - | - | - | 1723 | 2192 | 8639 |
| reduced edges | ER+ | - | - | - | 1322 | 2335 | 6031 |
| R<sup>2</sup> | both | - | - | - | 0.6 | 0.7 | 0.7 |
| WGCNA threshold range | both | - | - | - | 0.6-0.8 | 0.2-0.95 | 0.2-0.95 |

### 2 Additional analyses - Supplementary Figures

#### 2.1 Effect size for the ground truth

In the manuscript, we derived the ground truth by contrasting breast drug response measurements from multiple ER-stratified cancer cell lines from CTRP (Rees *et al*., 2015) with a Mann-Whitney-U test. We used the p-value as ground truth readout: the smaller the p-value, the more differential the drug response between conditions. Here, we investigated the impact of using the effect size instead of the p-value. The effect size is defined as 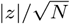, with *N* the sample size, i.e. the number of measured breast cancer cell lines for a drug, and *z* the z-score of the test statistic. We performed ROC analysis for DrDimont’s predictions, prediction by PageRank, or by differential protein expression for different thresholds on the effect size (see Figure S1). Overall, conclusions remain the same as for the original ground truth by p-values (compare Fig. 3 in the main text), with DrDimont performing better than the other two methods, and being predictive especially for top-ranked drugs.

**Fig. S1.**
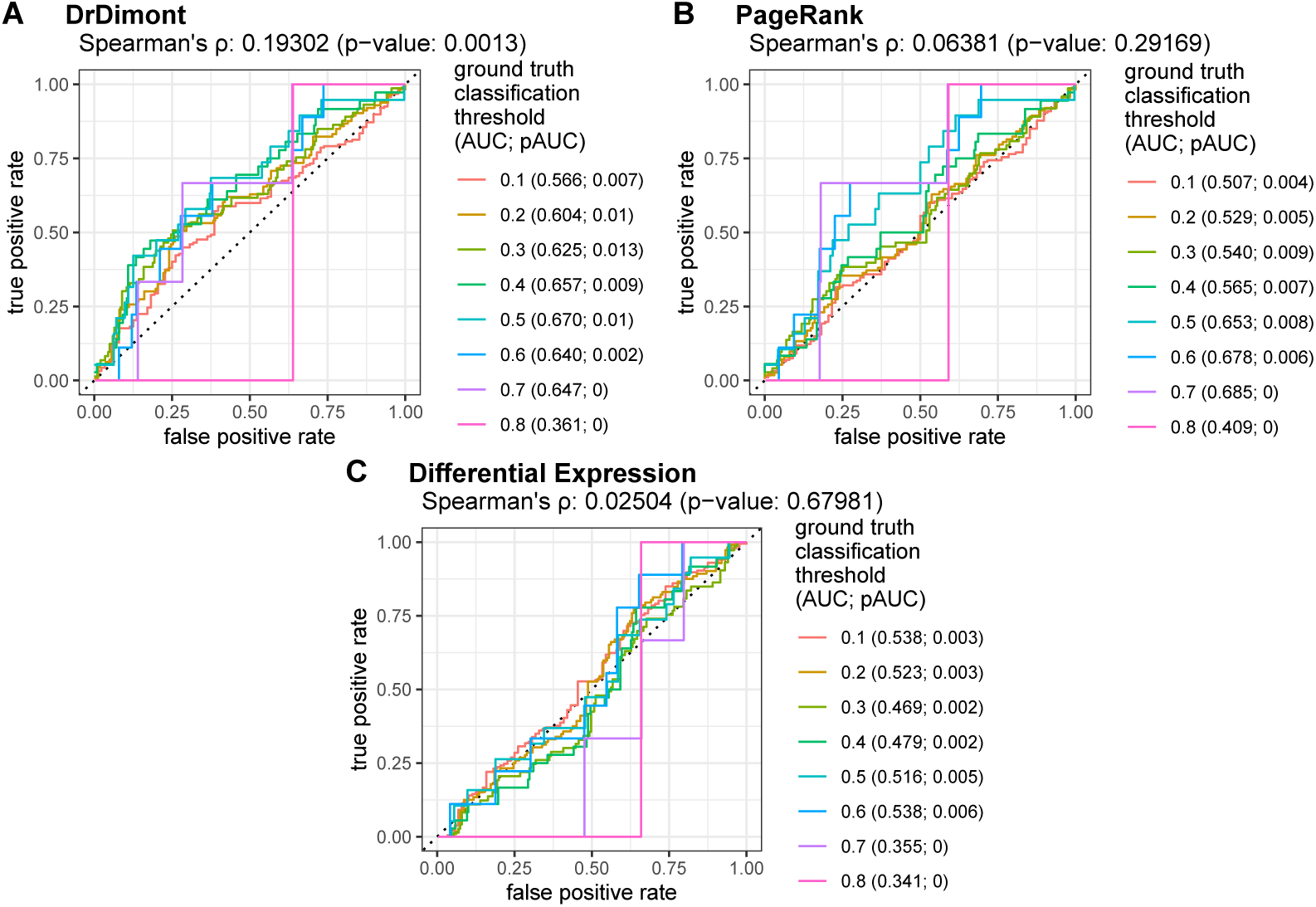
ROC curves for DrDimont’s and alternative methods’ prediction performance on the breast cancer dataset. Average effect size was used for ground truth drug ranking. (A) DrDimont’s differential drug response (275 drugs). (B) Weighted PageRank of drug targets for differential drug response prediction (275 drugs). (C) Differential expression of drug target proteins for differential drug response prediction (274 drugs). In the legends, the values in brackets denote AUCs and pAUCs at given ground truth thresholds. The respective Spearman’s *ρ* and the p-value are given at the top of each figure. Black dotted lines indicate theoretical ROC curves for random predictions.

#### 2.2 Alternative reduction methods

DrDimont performs a reduction of the single-layer networks (see Methods, *Single-layer network generation*). By default, networks are reduced to maximize their scale-freeness, setting a goodness-of-fit threshold, *R*^2^ (together with considerations on allowed network sizes). Other options implemented in DrDimont’s framework are reductions by an approximate network density or average edge count. This means that the weakest network edges are removed until a specific density,

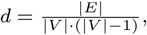

that is the ratio of existing edges to possible edges in the network, or a specific mean number of edges per node

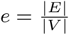

are reached. Here, *E* is the set of edges, *V* the set of vertices, and | *·* | denotes the cardinality of these sets. We investigated the effect of these alternative reduction schemes on DrDimont’s performance for the TCGA + CPTAC dataset (see Figure S2). We find that for certain network densities *d* and average numbers of edges per node *e*, prediction results are comparable to the *R*^2^-based reduction on scale-freeness. However, we observe that the prediction performance in terms of AUC and Spearman’s correlation to the ground truth decreases for sparse (*e* = 5, *d* = 0.001) networks, but also for very densely connected networks (*e* = 20, *d* = 0.007).

**Fig. S2.**
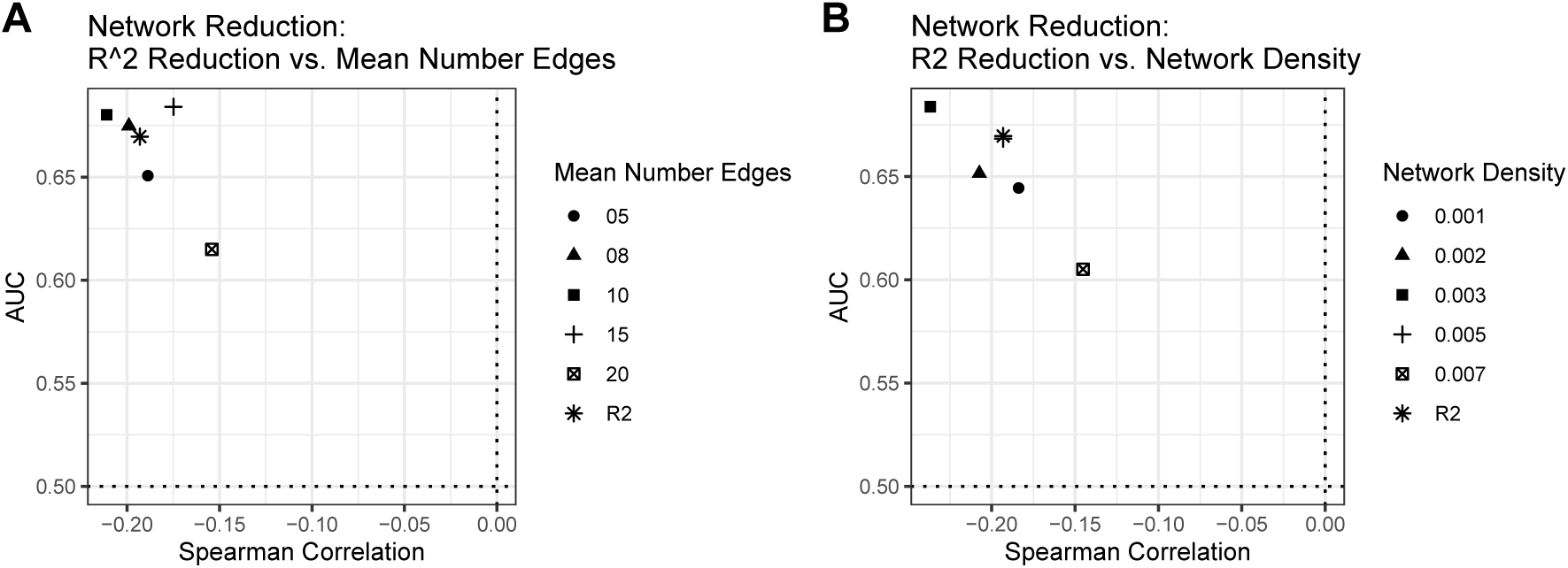
Effect of single-layer network reduction settings on DrDimont’s performance. AUCs vs. Spearman’s correlation for reduction upon (A) different average number of edges (5, 8, 10, 15, 20), (B) different network densities (0.001 - 0.007), compared to the prediction for reduction based on scale-freeness (R2). A ground truth threshold of 0.01 was used for AUC computation.

#### 2.3 Integrated interaction score settings and differential drug prediction method

By default, we propagate local edge information in the integrated interaction scores by considering alternative paths up to a length of *L* = 3. Larger path lengths are computationally difficult due to the extremely high number of possible paths. For smaller path lengths, we observe reduced performance, especially in terms of the correlation between prediction and ground truth (symbols in Figure S3, TCGA+CPTAC breast cancer dataset). This shows that the local merging scheme we devised is beneficial.

Furthermore, we examined the effect of summarizing the edge information around drug targets for drug prediction (see *Differential drug response* in the Methods of the main text). By default, we employ the arithmetic mean of edge weights in the differential network over all edges that are incident to the drug’s targets as differential drug response score of the drug (‘mean (difference)’). This results in the fact that strong interactions that are present only in one of the contrasted groups can be balanced by alternative strong interactions to other nodes in the other group. Alternatively, we here consider other measures: (i) the median (instead of mean) over those edge weights (‘median (difference)’), or (ii) absolute edge weights with mean (‘mean (absolute difference)’), or (iii) median (‘median (absolute difference)’) (colors in Figure S3). We find that our default measure performs decisively better than the other three, both in terms of AUC and correlation. Thereby, both measures that do not consider edge weights in absolute terms are better, and taking the mean is (at least slightly) better than taking the median.

**Fig. S3.**
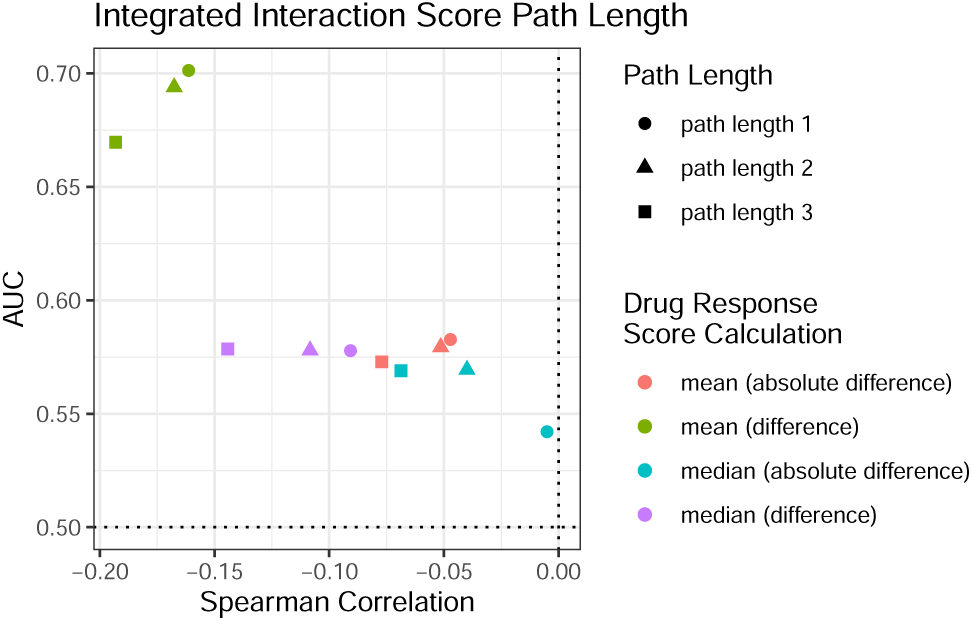
Integrated interaction score settings and differential drug computation. We examined DrDimont’s performance, AUC vs. correlation between prediction and ground truth (CTRP), on the TCGA + CPTAC dataset for different settings. We consider (i) the path length for computing the integrated interaction score: 3 (default, square), 2 (triangle), 1 (no propagation of local information, circle), and (ii) the summarizing scheme for edge weights incident to a drug’s targets in the differential network: mean of absolute differential edge weights (red), mean of differential edge weights (default, green), median of absolute differential edge weights (blue), and median of differential edge weights (purple).

#### 2.4 ROC curves for data layer ablation study

We here provide ROC curves for the data layer ablation study on the breast cancer patient dataset (from TCGA and CPTAC) with performance indicators from Table 1 from the main text. Default setting was to include the mRNA, protein, phosphosite layers (red in all plots in Figure S4). Apart from different combinations of these three layers, we assessed the impact of including data from one of two metabolomics studies (Budczies *et al*., 2013; Terunuma *et al*., 2014). Please refer to the main text for the description of the results.

**Fig. S4.**
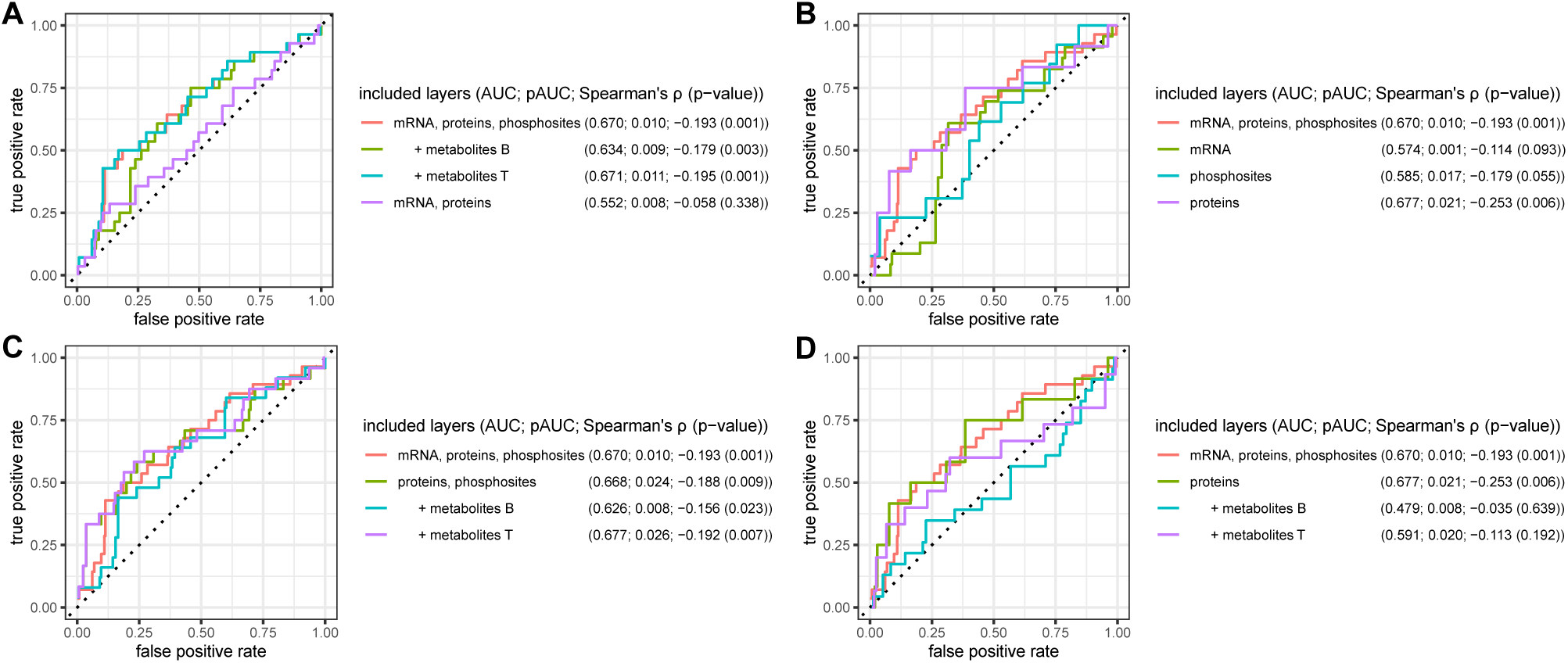
ROC curves for the performance of DrDimont when supplied different combinations of data layers for the TCGA + CPTAC breast cancer dataset. The default setting with mRNA, protein, phosphosite layers is given in each plot (red). (A) Addition of metabolomics, or removal of phosphosite data. (B) Single-layer prediction. (C) Protein and phosphosite layer, combined with metabolomics. (D) Protein layer, combined with metabolomics. Comparison against ground truth from CTRP; AUC and pAUC for a ground truth threshold of 0.01. B: Budczies dataset (Budczies et al., 2013), T: Terunuma dataset (Terunuma et al., 2014). Please refer to Table 1 (main text) for numbers of drugs with predictions in each setting.

#### 2.5 Drugs lacking predictions for data layers

We examined in more detail which drugs can be predicted as differential by DrDimont depending on the data layers that were provided. We focus on the case of using the protein layer only (best performing) or using the protein plus mRNA layers (strong decrease in performance). In particular, when using only the protein layer, we find that 159 of 275 drugs from CTRP that had drug targets in our molecular networks could not be predicted as differential or not by DrDimont. This happens if there are no edges incident to drug targets in either of the two compared conditions. Some of the most differential drugs (as indicated by CTRP ground truth) were among those (see Figure S5A). In contrast, when adding the mRNA layer, all 275 proteins receive DrDimont scores, but all but three of the ones without predictions for the protein layer only are zero (see Figure S5C). For drugs with DrDimont predictions in both settings, results are very similar (see Figure S5D).

The four drugs with highest differential predictions in both settings were AZ-3146, tamatinib, SB-743921, ibrutinib. The three drugs with high differential ground truth were AZ-D7762, MK1775, GSK461364. The drug with highest differential response in ground truth that received differential predictions in both settings was momelotinib (DrDimont’s drug response scores: 0.66 (protein), 0.63 (protein + mRNA)).

**Fig. S5.**
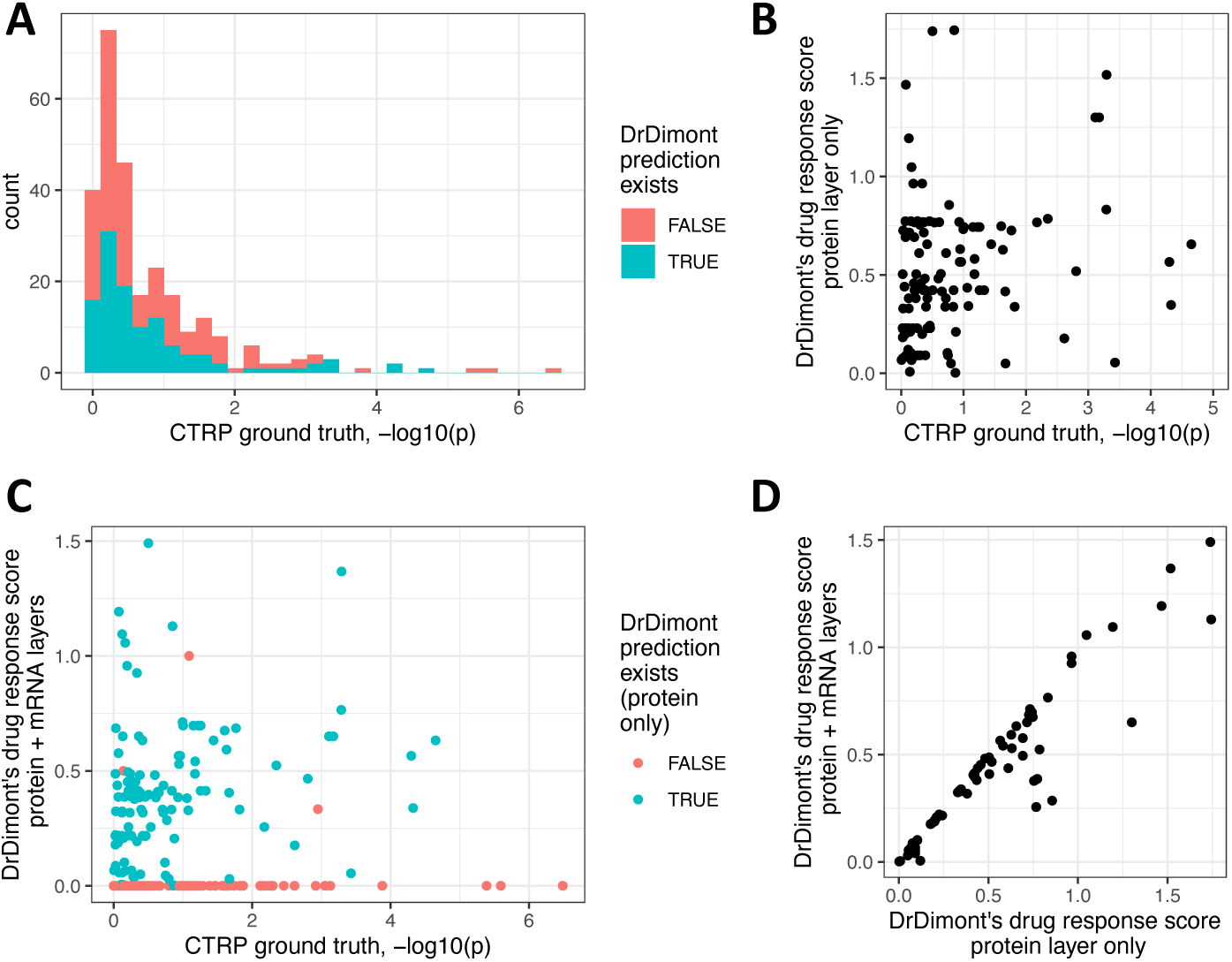
Effects of using only the protein layer or protein and mRNA layers on DrDimont’s predictions. (A) Histogram of ground truth predictions for CTRP (negative log10 of the Mann-Whitney-U p-values) for 275 drugs. The color distinguishes whether DrDimont could derive predictions for the drug from the protein layer only (116 drugs, blue) or not (159 drug, red). (B) DrDimont’s differential drug response score using only the protein layer vs. CTRP ground truth differential drug score predictions for 116 drugs. (C) DrDimont’s differential drug response score using the protein and mRNA layers vs. CTRP ground truth differential drug score predictions for all 275 drugs. Color indicates whether DrDimont could derive predictions from the protein layer only (as in A). (D) DrDimont’s differential drug response scores using the protein layer and mRNA layers vs. using only the protein layer for 116 drugs.

#### 2.6 Alternative ground truth measurements from GDSC

In our analyses, we focused on CTRP (Rees *et al*., 2015) as differential drug response ground truth because it delivered values for many drugs (490 measured, 275 with predictions in our framework). In order to assess the impact of this selected ground truth, we also employed the processed GDSC drug sensitivity dataset (Yang *et al*., 2013) that delivers data for 42 breast cancer cell lines (29 ER-, 13 ER+) and 198 drugs. From that, we used the area under the drug response curves (AUC) as drug sensitivity information. We retrieved the drug target information from the compound annotation file and matched protein names or target descriptions to Gene Symbols using the GeneCards resource. We only employed data for a drug if it was measured at least three times for each ER+ and ER-cell lines. We determined the differential drug response between ER+ and ER-for each drug by a Mann-Whitney-U test comparing sensitivity in ER+ cell lines vs. sensitivity in ER-cell lines. Thus, we obtained ground truth for 190 drugs from GDSC. For the dataset with TCGA mRNA, CPTAC proteomics and phosphosites, DrDimont predicted differential response for 101 drugs. A comparison between the ground truth from GDSC and CTRP and performance of DrDimont for them is shown in Figure S6. It becomes clear that the ground truths do not overlap very well.

**Fig. S6.**
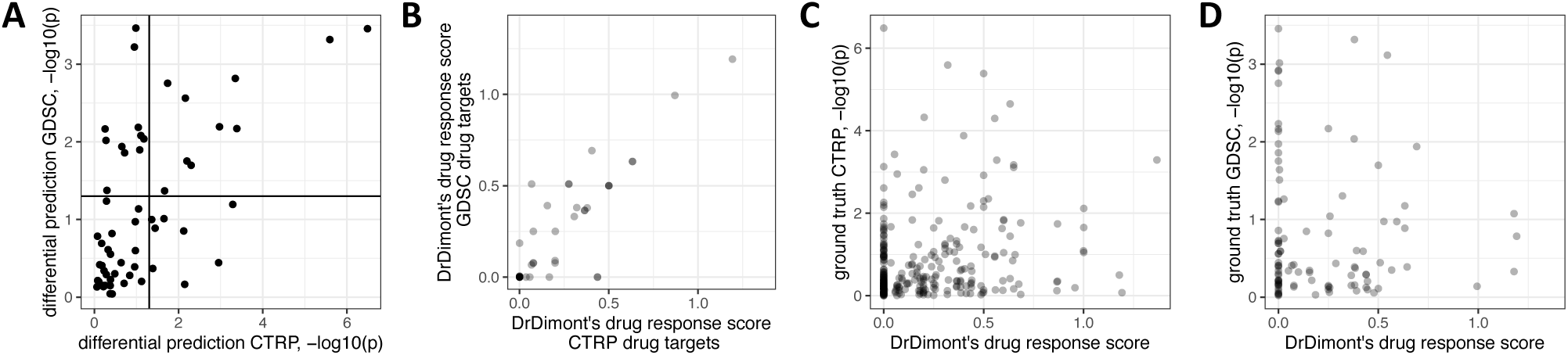
CTRP vs. GDSC ground truth for differential drug response. (A) Differential response ground truth negative logarithmized p-values for 57 drugs that overlap in the datasets from CTRP and GDSC. Solid lines indicate p-values of 0.05. (B) DrDimont’s drug response scores for the TCGA + CPTAC breast cancer dataset (mRNA, protein, phosphosites) for 53 drugs overlapping between CTRP and GDSC. Different scores stem from different drug target definitions, 4 of the 57 drugs from (A) did not have drug targets in our network. (C) DrDimont’s drug response scores compared to CTRP ground truth for 275 drugs. Dataset as in B. Performance at a 0.01 ground-truth threshold: AUC 0.67, pAUC 0.01; Spearman’s *ρ* -0.193, p-value 0.001. (D) DrDimont’s drug response scores compared to GDSC ground truth for 101 drugs. Dataset as in B. Performance at a 0.01 ground-truth threshold: AUC 0.4, pAUC 0; Spearman’s *ρ* 0.01, p-value 0.9.

#### 2.7 Performance for unreduced and other datasets

In addition to the TCGA + CPTAC dataset, we applied DrDimont to two other datasets for ER+ vs. ER-breast cancer: (i) a second independent breast cancer patient study with mRNA, protein, phosphosites (Krug *et al*., 2020) (‘Krug’), and (ii) cell line data from the Cancer Cell Line Encyclopedia (CCLE) with mRNA, proteins and metabolomics (Li *et al*., 2019; Nusinow *et al*., 2020). The mRNA, protein and phosphosite data for the Krug dataset was derived from the Supplementary Table 2 of Krug et al. (Krug *et al*., 2020). The breast cancer cell line data was downloaded from the CCLE (Iorio *et al*., 2016; Nusinow *et al*., 2020; Li *et al*., 2019).

For the Krug dataset, we only employed samples with tumor purity (according to the metadata) > 0.5; the ER status of the samples was reported in the metadata file (Krug *et al*., 2020). For the cell line data, we used estrogen receptor status annotation from the CCLE resource. Both mRNA datasets stem from RNAseq. We neglected mRNAs with more than 90% of zero measurements over the samples within a condition. Proteins and phosphosites with more than 20% of missing values over the samples of a condition were removed. DrDimont’s performance was compared to predictions derived from differential protein expression analysis (see Methods, *Differential protein expression*).In addition, we re-analyzed the TCGA + CPTAC (‘TCGA’) dataset without genetic feature reduction, and no genetic feature selection was performed for the other two datasets either. See Table S1 for DrDimont’s settings, sample sizes for each subgroup of the datasets, and generated network sizes.

**Fig. S7.**
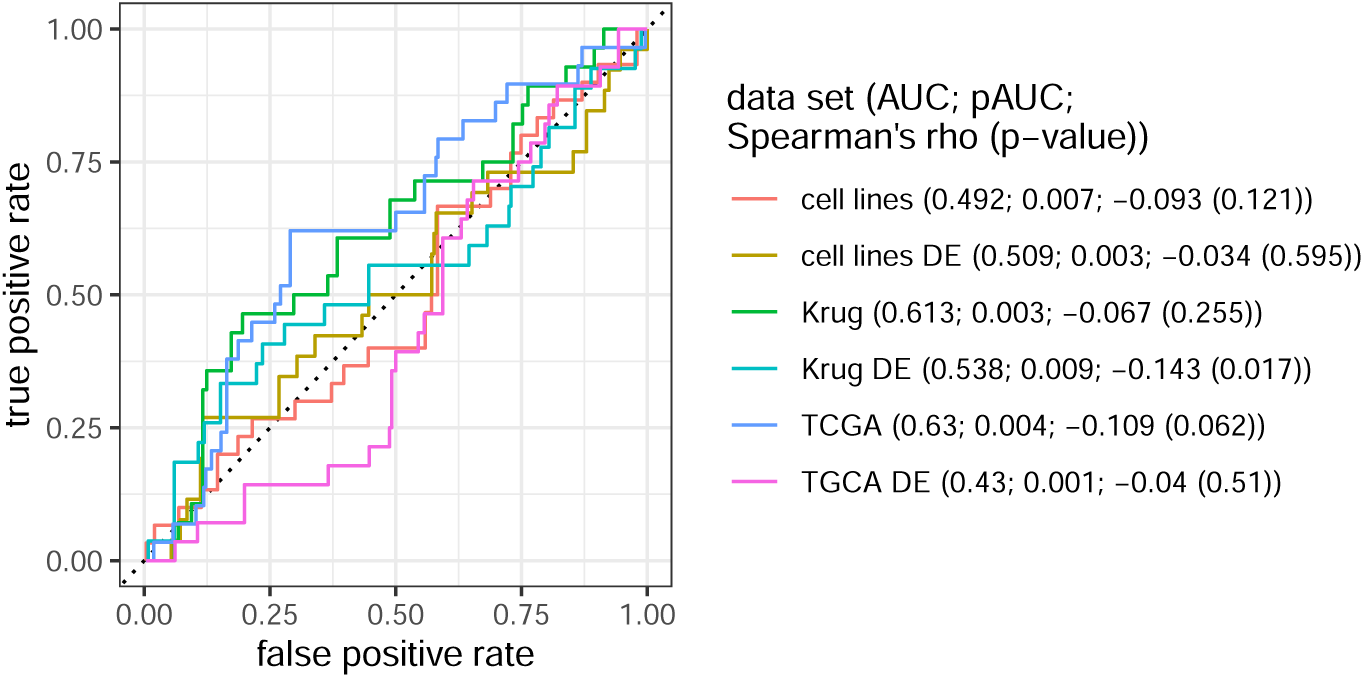
DrDimont’s differential drug prediction results for three different datasets, compared to prediction by differential expression (‘DE’): breast cancer cell lines (‘cell lines’), an alternative breast cancer patient study (‘Krug’), and TCGA + CPTAC breast cancer patients (‘TCGA’). Genetic features were not reduced. ROC analysis was performed with a ground-truth threshold of 0.01 for the CTRP ground truth. The AUC, pAUC and Spearman’s *ρ* and its significance (p-value) are described in the legend. Predictions are derived for 278 drugs in the cell lines dataset, 295 drugs in the Krug dataset, and 292 drugs in the entire TCGA dataset.

For the cell line data, DrDimont’s prediction showed a performance close to random in terms of AUC (0.492), but elevated pAUC (0.007 vs. 0.005 expected for random predictions) and also a small correlation to the ground truth prediction (Spearman’s *ρ* -0.093, p-value 0.12). The drug response prediction by differential expression performed worse with even lower correlation (-0.034) and decisively lower pAUC (0.003). Reasons for the bad performance on the cell line dataset could be the decisively lower sample size that characterizes each subgroup. In addition, we lack the phosphosite layer for this dataset that had shown to confer robustness to the results in our TCGA + CPTAC dataset.

For the Krug dataset, DrDimont’s predictions yielded an AUC of 0.61 and an insignificant Spearman’s *ρ* of -0.07 (p-value 0.255). Top-ranked results were not retrieved particularly frequently (pAUC 0.003). For differential drug response by differential protein expression in the Krug dataset, lower AUC (0.538), but significant correlation (-0.14, p-value 0.017) and a higher pAUC than expected by random predictions (0.009) were obtained. Therefore, in this dataset, it seems that differential protein expression is a better marker for differential drug response than network-based interaction analysis. This suggests that including differential expression into the analysis approaches might be beneficial under certain circumstances. DrDimont’s drug response scores of the unreduced TCGA dataset resulted in slightly worse predictions than for the reduced dataset (AUC 0.63 for ground-truth threshold 0.01, Spearman’s *ρ* -0.11, p-value 0.062), and especially lacked enrichment of positive predictions for top-ranked results (pAUC 0.004). Thus, feature selection seems beneficial for differential drug response.

## References

Allaire, J., et al. (2022). rmarkdown: Dynamic documents for R. https://github.com/rstudio/rmarkdown.

Argelaguet, R., et al. (2018). Multi-omics factor analysis-a framework for unsupervised integration of multi-omics data sets. Mol Syst Biol, 14(6), e8124.

Attali, D. and Baker, C. (2022). ggExtra: Add marginal histograms to ‘ggplot2’, and more ‘ggplot2’ enhancements. https://CRAN.R-project.org/package=ggExtra.

Azuaje, F. (2019). Artificial intelligence for precision oncology: beyond patient stratification. npj Precision Oncology, 3(1), 6.

Bache, S. M. and Wickham, H. (2022). magrittr: A forward-pipe operator for R. https://CRAN.R-project.org/package=magrittr.

Barretina, J., et al. (2012). The Cancer Cell Line Encyclopedia enables predictive modelling of anticancer drug sensitivity. Nature, 483(7391), 603–607.

Bartel, J., et al. (2015). The human blood metabolome-transcriptome interface. PLOS Genetics, 11(6), e1005274.

Bersanelli, M., et al. (2016). Methods for the integration of multi-omics data: mathematical aspects. BMC Bioinformatics, 17 Suppl 2, 15.

Bhuva, D. D., et al. (2019). Differential co-expression-based detection of conditional relationships in transcriptional data: comparative analysis and application to breast cancer. Genome Biology, 20(1), 236.

Brin, S. and Page, L. (1998). The anatomy of a large-scale hypertextual web search engine. Computer Networks and ISDN Systems, 30(1), 107–117.

Budczies, J., et al. (2013). Comparative metabolomics of estrogen receptor positive and estrogen receptor negative breast cancer: alterations in glutamine and beta-alanine metabolism. J Proteomics, 94, 279–88.

Cantini, L., et al. (2021). Benchmarking joint multi-omics dimensionality reduction approaches for the study of cancer. Nature Communications, 12(1), 124.

Chang, K., et al. (2013). The Cancer Genome Atlas Pan-Cancer analysis project. Nature Genetics, 45(10), 1113–1120.

Csardi, G. and Nepusz, T. (2006). The igraph software package for complex network research. InterJournal, Complex Systems, 1695.

da Costa-Luis, C., et al. (2021). tqdm: A fast, extensible progress bar for Python and CLI. https://doi.org/10.5281/zenodo.5202772.

Demirel, H. C., et al. (2021). Computational approaches leveraging integrated connections of multi-omic data toward clinical applications. Molecular Omics.

DepMap, Broad (2021). DepMap 21Q4. Public figshare dataset. https://doi.org/10.6084/m9.figshare.16924132.v1.

Di Nanni, N., et al. (2020). Network diffusion promotes the integrative analysis of multiple omics. Frontiers in Genetics, 11, 106.

Dimitrakopoulos, C., et al. (2018). Network-based integration of multi-omics data for prioritizing cancer genes. Bioinformatics, 34(14), 2441–2448.

Ding, M. Q., et al. (2018). Precision oncology beyond targeted therapy: Combining omics data with machine learning matches the majority of cancer cells to effective therapeutics. Mol Cancer Res, 16(2), 269–278.

Dittmar, G., et al. (2019). PRISMA: Protein interaction screen on peptide matrix reveals interaction footprints and modifications-dependent interactome of intrinsically disordered C/EBPbeta. iScience, 13, 351–370.

Gao, H., et al. (2015). High-throughput screening using patient-derived tumor xenografts to predict clinical trial drug response. Nature Medicine, 21(11), 1318–1325.

Geeleher, P., et al. (2017). Discovering novel pharmacogenomic biomarkers by imputing drug response in cancer patients from large genomics studies. Genome Res, 27(10), 1743–1751.

Harrell Jr, F. (2021). Hmisc: Harrell miscellaneous. https://CRAN.R-project.org/package=Hmisc.

Harris, C. R., et al. (2020). Array programming with NumPy. Nature, 585(7825), 357–362.

Henry, L. and Wickham, H. (2022). rlang: Functions for base types and core R and ‘tidyverse’ features. https://CRAN.R-project.org/package=rlang.

Huang, S., et al. (2017). More is better: Recent progress in multi-omics data integration methods. Front Genet, 8, 84.

Ideker, T. and Krogan, N. J. (2012). Differential network biology. Molecular Systems Biology, 8(1), 565.

Iorio, F., et al. (2016). A landscape of pharmacogenomic interactions in cancer. Cell, 166(3), 740–754.

Koh, H. W. L., et al. (2019). iOmicsPASS: network-based integration of multiomics data for predictive subnetwork discovery. NPJ Syst Biol Appl, 5, 22.

Kosinski, M. and Biecek, P. (2021). RTCGA: The Cancer Genome Atlas Data Integration. https://rtcga.github.io/rtcga.

Krug, K., et al. (2020). Proteogenomic landscape of breast cancer tumorigenesis and targeted therapy. Cell, 183(5), 1436–1456.e31.

Langfelder, P. and Horvath, S. (2008). WGCNA: an R package for weighted correlation network analysis. BMC Bioinformatics, 9, 559.

Lee, B., et al. (2019). Heterogeneous multi-layered network model for omics data integration and analysis. Front Genet, 10, 1381.

Leucht, S., et al. (2015). How effective are common medications: a perspective based on meta-analyses of major drugs. BMC Medicine, 13(1), 253.

Li, H., et al. (2019). The landscape of cancer cell line metabolism. Nat Med, 25(5), 850–860.

Matsui, Y., et al. (2021). RoDiCE: robust differential protein co-expression analysis for cancer complexome. Bioinformatics.

Mertins, P., et al. (2016). Proteogenomics connects somatic mutations to signalling in breast cancer. Nature, 534(7605), 55–62.

Moritz, P., et al. (2018). Ray: A distributed framework for emerging AI applications. In Proceedings of the 13th USENIX Conference on Operating Systems Design and Implementation, OSDI’18, page 561–577, USA. USENIX Association.

Müller, K. and Wickham, H. (2021). tibble: Simple data frames. https://CRAN.R-project.org/package=tibble.

Nusinow, D. P., et al. (2020). Quantitative proteomics of the Cancer Cell Line Encyclopedia. Cell, 180(2), 387–402.e16.

Ogris, C., et al. (2021). Versatile knowledge guided network inference method for prioritizing key regulatory factors in multi-omics data. Scientific Reports, 11(1), 6806.

Ortmayr, K., et al. (2019). Metabolic profiling of cancer cells reveals genome-wide crosstalk between transcriptional regulators and metabolism. Nature Communications, 10(1), 1841.

Papadakis, M., et al. (2022). Rfast: A collection of efficient and extremely fast R functions. https://CRAN.R-project.org/package=Rfast.

Park, A., et al. (2022). A comprehensive evaluation of regression-based drug responsiveness prediction models, using cell viability inhibitory concentrations (IC50 values). Bioinformatics, page btac177.

Park, Y. H., et al. (2020). Chemotherapy induces dynamic immune responses in breast cancers that impact treatment outcome. Nature Communications, 11(1), 6175.

Parvizi, P., et al. (2020). A network-based embedding method for drug-target interaction prediction. Annu Int Conf IEEE Eng Med Biol Soc, 2020, 5304–5307.

Peng, C., et al. (2017). Discovery of bladder cancer-related genes using integrative heterogeneous network modeling of multi-omics data. Scientific Reports, 7(1), 15639.

Picard, M., et al. (2021). Integration strategies of multi-omics data for machine learning analysis. Computational and Structural Biotechnology Journal, 19, 3735–3746.

Pinto, J. P., et al. (2014). Targeting molecular networks for drug research. Frontiers in Genetics, 5.

Pinu, R. F., et al. (2019). Systems biology and multi-omics integration: Viewpoints from the metabolomics research community. Metabolites, 9(4).

Pio-Lopez, L., et al. (2021). Multiverse: a multiplex and multiplex-heterogeneous network embedding approach. Scientific Reports, 11(1), 8794.

Python Core Team (2021). Python: A dynamic, open source programming language. https://www.python.org/.

R Core Team (2021). R: A language and environment for statistical computing. https://www.r-project.org/.

Recanatini, M. and Cabrelle, C. (2020). Drug research meets network science: Where are we? Journal of Medicinal Chemistry, 63(16), 8653–8666.

Rees, M. G., et al. (2015). Correlating chemical sensitivity and basal gene expression reveals mechanism of action. Nature Chemical Biology, 12, 109.

Repana, D., et al. (2019). The Network of Cancer Genes (NCG): a comprehensive catalogue of known and candidate cancer genes from cancer sequencing screens. Genome Biology, 20(1), 1.

Richard, M., et al. (2020). PenDA, a rank-based method for personalized differential analysis: Application to lung cancer. PLOS Computational Biology, 16(5), e1007869.

Sambaturu, N., et al. (2020). PathExt: a general framework for path-based mining of omics-integrated biological networks. Bioinformatics.

Schulte-Sasse, R., et al. (2021). Integration of multiomics data with graph convolutional networks to identify new cancer genes and their associated molecular mechanisms. Nature Machine Intelligence.

Sompairac, N., et al. (2019). Independent component analysis for unraveling the complexity of cancer omics datasets. Int J Mol Sci, 20(18).

Stanfield, Z., et al. (2017). Drug response prediction as a link prediction problem. Scientific Reports, 7(1), 40321.

Szklarczyk, D., et al. (2016). STITCH 5: augmenting protein-chemical interaction networks with tissue and affinity data. Nucleic Acids Res, 44(D1), D380–4.

Terunuma, A., et al. (2014). Myc-driven accumulation of 2-hydroxyglutarate is associated with breast cancer prognosis. J Clin Invest, 124(1), 398–412.

Valdeolivas, A., et al. (2019). Random walk with restart on multiplex and heterogeneous biological networks. Bioinformatics, 35(3), 497–505.

Wang, B., et al. (2014). Similarity network fusion for aggregating data types on a genomic scale. Nat Methods, 11(3), 333–7.

Webber, J. T., et al. (2018). Integration of tumor genomic data with cell lines using multi-dimensional network modules improves cancer pharmacogenomics. Cell Syst, 7(5), 526–536 e6.

Wickham, H. (2016). ggplot2: Elegant Graphics for Data Analysis. Springer-Verlag New York.

Wickham, H. (2019). stringr: Simple, consistent wrappers for common string operations. https://CRAN.R-project.org/package=stringr.

Wickham, H. and Girlich, M. (2022). tidyr: Tidy messy data. https://CRAN.R-project.org/package=tidyr.

Wickham, H., et al. (2021). dplyr: A grammar of data manipulation. https://CRAN.R-project.org/package=dplyr.

Wickham, H., et al. (2022). readr: Read rectangular text data. https://CRAN.R-project.org/package=readr.

Wishart, D. S., et al. (2017). Drugbank 5.0: a major update to the drugbank database for 2018. Nucleic Acids Research, 46(D1), D1074–D1082.

Xie, Y. (2022). knitr: A general-purpose package for dynamic report generation in R. https://yihui.org/knitr/.

Yang, W., et al. (2013). Genomics of Drug Sensitivity in Cancer (GDSC): a resource for therapeutic biomarker discovery in cancer cells. Nucleic Acids Research, 41(D1), D955–D961.

Yugi, K., et al. (2016). Trans-omics: How to reconstruct biochemical networks across multiple ‘omic’ layers. Trends Biotechnol, 34(4), 276–90.

Zhang, F., et al. (2018a). A novel heterogeneous network-based method for drug response prediction in cancer cell lines. Scientific Reports, 8(1), 3355.

Zhang, X., et al. (2018b). Proteome-wide identification of ubiquitin interactions using UbIA-MS. Nat Protoc, 13(3), 530–550.

Zhu, L., et al. (2016). KSP inhibitor SB743921 inhibits growth and induces apoptosis of breast cancer cells by regulating p53, Bcl-2, and DTL. Anticancer Drugs, 27(9), 863–72.

